# Aberrant circuitry underlying olfaction in the face of severe olfactory bulb degeneration

**DOI:** 10.1101/2023.01.31.526422

**Authors:** Tamar Licht, Michael Yunerman, Ido Maor, Naheel Lawabny, Renana Oz Rokach, Adi Mizrahi, Dan Rokni

**Affiliations:** Department of Medical Neurobiology, Faculty of Medicine and IMRIC, The Hebrew University of Jerusalem, Jerusalem, Israel; Department of Neurobiology, The Edmond and Lily Safra Center for Brain Sciences, The Hebrew University of Jerusalem, Jerusalem, Israel

**Author notes:** Corresponding authors. Dan Rokni, Tamar Licht. These authors contributed equally to this work (Tamar Licht, Michael Yunerman).

## Abstract

The olfactory bulb (OB) is a critical component of mammalian olfactory neuroanatomy. Beyond being the first and sole relay station for olfactory information to the rest of the brain, it also contains elaborate stereotypical circuitry that is considered essential for olfaction. Indeed, substantial lesions of the OB in rodents lead to anosmia. Here, we examined the circuitry that underlies olfaction in a mouse model with severe developmental degeneration of the OB. These mice could perform odor-guided tasks and even responded normally to innate olfactory cues. Despite the near total loss of the OB, piriform cortex in these mice responded to odors normally and its neural activity sufficed to decode odor identity. We analyzed the circuitry that supports olfactory function in these mice. We found that sensory neurons express the full repertoire of olfactory receptors and their axons project primarily to the rudimentary OB, but also ectopically, to olfactory cortical regions. Within the OB, the number of principal neurons was greatly reduced and the morphology of their dendrites was abnormal, extending over larger regions within the OB. Glomerular organization was lost. This study shows that olfactory functionality can be preserved despite reduced and aberrant circuitry that is missing many of the elements that are believed to be essential for olfaction, and may explain the retention of olfaction in humans with degenerated OBs.

## Introduction

Odor sensing begins with the specific binding of odorants to receptors that are expressed on the membranes of olfactory sensory neurons (OSNs) in the nasal epithelium. OSNs project to the olfactory bulb (OB) that performs basic processing functions and projects further to several brain regions that are collectively regarded as olfactory cortex. The OB is therefore considered essential for olfaction not only because it provides the sole link between the nose and the brain, but also because it is assumed to perform unique computations (Mori et al., 1999; Schoppa and Urban, 2003; Wachowiak and Shipley, 2006; Wilson and Mainen, 2006; Linster and Cleland, 2009). Studies in rodents indeed found that olfaction is lost if OB lesions are extensive (Lu and Slotnick, 1998; Erskine et al., 2019). It was therefore surprising that some humans maintain a functional sense of smell despite the absence of apparent OBs (Rombaux et al., 2007; Weiss et al., 2020). Plausible explanations for this finding may be that a small degenerated OB suffices for a functional sense of smell, or, that the loss of the OB early in development is compensated for if sensory neurons manage to find novel synaptic targets (Slotnick et al., 2004). The young brain is known to be plastic and capable of regeneration, which allows it to compensate for damage. Loss of function of one part of the brain is often compensated for by other parts that modify their connectivity and acquire new functions. This was described in both motor(Jones, 2017) and sensory systems (Slotnick et al., 2004; Depner et al., 2014; Takiguchi et al., 2015). The olfactory system is particularly plastic, having niches for adult neurogenesis in both the OB and the olfactory epithelium (OE) (Schwob et al., 1995; Imitola et al., 2004). Constant replacement of OSNs enables the olfactory system to continuously adapt to environmental changes (Ibarra-Soria et al., 2017; Tsukahara et al., 2021) as well as restore function following damage (Schwob et al., 1995). The extreme plasticity of the olfactory system may therefore enable it to preserve its function despite developmental abnormalities (Fleischmann et al., 2008; Knott et al., 2012; Roland et al., 2016).

Here, we used a mouse model for severe developmental OB degeneration to reveal how olfaction may be maintained in subjects with degenerated OBs. This model, based on OB-specific blood vessel regression during development, generates healthy adults in which the OB volume is reduced to as little as 3% of the normal size. We assessed innate and learned olfactory skills, recorded odor responses in piriform cortex and studied the various abnormalities and adaptations at the histological/cellular level.

## Results

### A genetic system for OB degeneration during development

We used a mouse model with severe developmental OB degeneration to study how olfaction is maintained despite the extreme anatomical aberration. In these mice, inducible, brain-specific expression of the human VEGF-binding domain of VEGFR1 (sFlt1), sequesters VEGF and blocks its angiogenic signaling. Blood vessel growth and maintenance rely on VEGF at early stages but become refractory to VEGF upon maturation (Benjamin et al., 1999; Licht et al., 2015). Blocking VEGF signaling prior to blood vessel maturation, leads to a collapse of the blood vessels and degeneration of the surrounding neural tissue. We exploited the fact that OB vasculature is the last to mature in the brain and blocked VEGF signaling at E13.5 to create mice with severe degeneration of the OB (Figure 1) (Licht et al., 2010, 2015). The size of the remaining OB varied from 3% to 60% of its normal size (Figure 1d, OB volumes are presented as % of the mean control OB volume - 10.05±0.36, mean±SEM, n=6), while other parts of the brain were little affected (Supp Figure 1). In the rest of this manuscript, we analyze olfactory function and the underlying circuitry in these mice (mice in this study and their respective OB volume appear in Supplementary Table 1).

**Figure 1.**
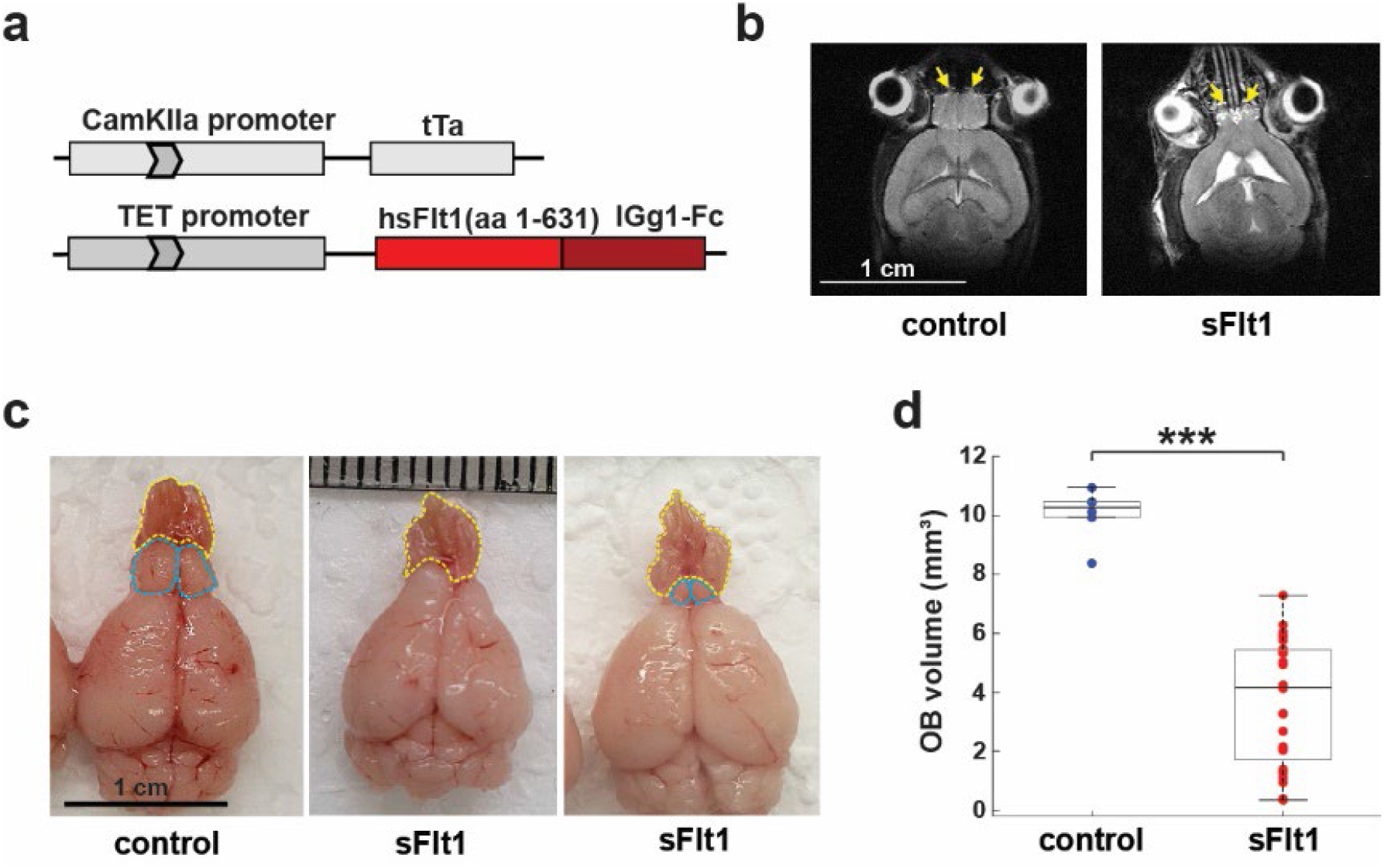
An experimental system for conditional VEGF loss-of-function leading to OB degeneration. (**a**) Diagram of the ‘driving’ CamkIIα promoter - tTA transgene and the ‘responder’ tetracycline-regulated sFlt1 transgene composing the conditionally-induced bi-transgenic system for VEGF blockade. (**b**) MRI scans of one control and one sFlt1 mouse (axial plane) for estimating OB size prior to behavioral experiments. Arrows indicate OBs. (**c**) Brains from one control and two sFlt1 mice aged 5 months. The OB and OE are labeled in blue and yellow, respectively. (**d**) OB volumes of 6 control mice and 28 sFlt1 mice, calculated according to axial plane projections of MRI scans P=3.34E^−10^ (t-test).

### sFlt1 mice can perform odor-guided behaviors

We first examined whether sFlt1 mice with degenerated OBs maintained a functioning sense of smell. Notably, pups could nurse from their mothers as they develop to be adults gaining normal body weight (>20 gr) and could also mate and produce offspring.

We started with the buried food test in which the speed of localizing hidden food is taken as a measure of olfactory ability (Yang and Crawley, 2009). Mice were placed in an arena with a food pellet hidden under the bedding and the time to find the pellet was measured. Each mouse performed the task 4 times and the average search time was calculated. All mice completed the task in less than 5 minutes with no significant difference between the control and sFlt1 groups, indicating that sFlt1 mice can detect and localize odors sources (Figure 2a). We next asked whether sFlt1 mice retain the normal innate valence of various odors. To probe valence, we measured the attraction/repulsion of mice to several natural odors with clear ethological significance for mice (female mouse urine, male rat urine, and 2,4,5-trimethyl-3-thiazoline (TMT, a well-known aversive odorant(Ayers et al., 2013)). We placed mice in an arena that contained the odor-source in one of its corners and monitored their movement for 5 minutes (Figure 2b). The percent of time a mouse spent in the quadrant of the arena that contained the odor-source, was used as a measure of odor valence. All 3 odors affected place preference with no significant difference between sFlt1 mice and controls (Figure 2c-e). Mice were attracted to mouse urine and surprisingly also to rat urine, and were repelled by TMT. Importantly, the performance of individual mice on the self-motivated tasks was not correlated with OB size (Supp Figure 2a-d, p=0.32, p=0.78, and p=0.53 for female urine, rat urine, and TMT, respectively), indicating that these tasks either require very little OB-dependent processing or that they were rescued by compensatory circuitry. Placing the mice in the same arena without any odor-source revealed no preference for a specific quadrant (Figure 2f).

**Figure 2.**
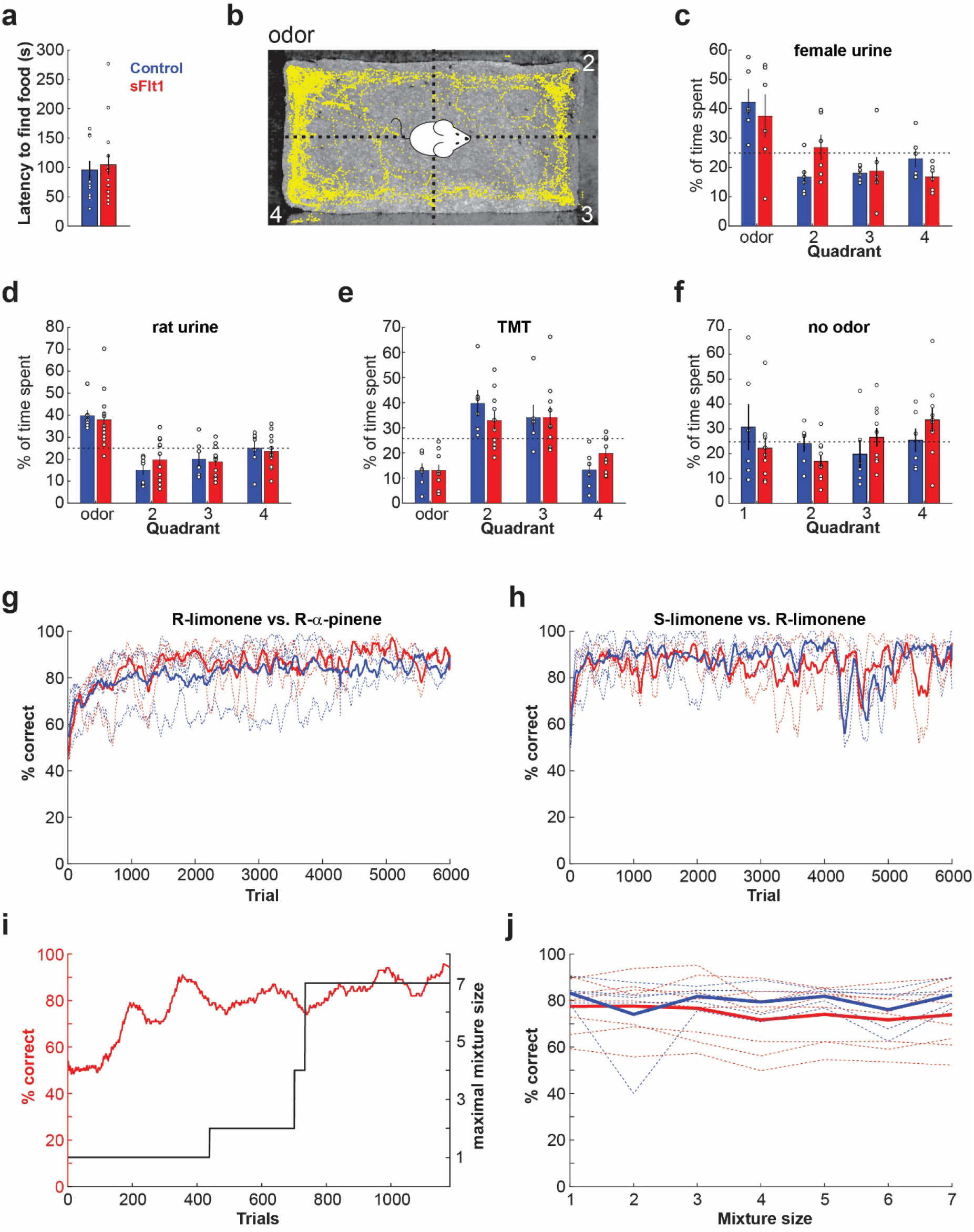
Behavioral performance on odor-guided tasks. (**a**) The latency to find a hidden food pellet buried 3cm under the bedding. Each dot is the average of 4 sessions per animal (sessions held once a week after 18h of food restriction). N=11 controls, 14 sFlt1, males and females. p = 0.71, Wilcoxon rank sum. Blue and red in all panels denote control and sFlt1 mice, respectively. (**b**) Open-field odor-guided place preference. Odors were smeared in a corner of the arena. Mice were placed in the center and video-tracked for 5 min. The percentage of time spent in each quadrant was calculated. (**c**) Freshly-collected female urine (mix of 5-6 females). Only male mice were tested (n=6 controls and 6 sFlt1). Main effect (2-way ANOVA): significant interaction of place F(3,30)=8.86 p=2.33*10^−4^, insignificant interaction of group (P=0.328). Post hoc test showed a significant difference between the urine quadrant and the other 3 quadrants in the control group (P=0.004, 0.039, 0.002) and in the sFlt1 group (P=0.048, 0.02, 0.002). (**d**) Freshly-collected male rat urine. Males and females were tested (N=7 controls and 12 sFlt1). Main effect (2-way ANOVA): significant interaction of place F(3,51)=15.827 p=2.11*10^−7^, insignificant interaction of group (p=0.706). Post hoc test showed a significant difference between the urine quadrant and the other 3 quadrants in the control group (P=5.03*10^−5^, 0.05, 0.002) and in the sFlt1 group (P=9.35*10^−5^, 0.004, 9.35*10^−5^). (**e**) TMT, 10μl. Males and females were tested (N=6 controls and 10 sFlt1). Main effect (2-way ANOVA): significant interaction of place F(3,42)=14.95 p=9.1*10^−7^, insignificant interaction of group (p=0.52). Post hoc test showed a significant difference between the TMT quadrant and the opposite 2 quadrants in the control group (P=0.001, 0.001) and in the sFlt1 group (P=0.001, 0.02). (**f**) Place preference with no odor added to the arena (N=6 controls and 10 sFlt1) showed no significant interactions. place: p=0.195, group: p=0.169. (**g-h**) Learning curves of control and sFlt1 mice in the Educage odor discrimination go/nogo task. Mice were first trained to discriminate between R-limonene (Go) and pinene (no-go) (**g**), and then between S-limonene (go) vs. R-limonene (no-go) (**h**). N=4 controls (2 males, 2 females), 3 sFlt1 (2 males, 1 female). OB volumes 10-30% of normal. The OB of one of the sFlt1 mice is presented in Figure 5a. Dashed lines show the performance of individual mice; thick lines are the group averages. (**i-j**) Olfactory figure-background segmentation. (**i**) Red, learning curve of a representative sFlt1 mouse (OB size 2mm^3^, about 20% of the normal size, images of this mouse’s OB appear in Figure 6g). Black, the maximal mixture size in the session. (**j**) performance of 7 sFlt1 mice (3 females, 4 males) and 6 controls, as a function of mixture size.

To probe olfactory acuity in sFlt1 mice, we next tested structured odor-guided tasks in which mice are trained to respond to odors according to predetermined rules and in limited time. We started with a simple 2-odor discrimination task using an automated training system called the Educage (Maor et al., 2019). In this setup, mice are free to engage a behavioral port at their own will, where they consume their entire water intake. Following a habituation period, mice were trained to perform a go/no-go olfactory discrimination task in which they were required to lick in response to one odor and refrain from licking in response to another. Mice were initially trained to discriminate R-limonene vs R-pinene. All mice reached performance levels of >80% with no significant difference between sFlt1 mice and controls (Figure 2g). Limonene and pinene have very distinct smells (lemon and pine, respectively), and are easy to discriminate. To further increase the difficulty of the task, we trained the same mice to discriminate between two enantiomers: R- vs S- limonene (Blount and Coppola, 2020). Mice were similarly successful in this task achieving performance levels of >80%, again with no significant differences between controls and sFlt mice (Figure 2h).

Beyond discriminating odors, the olfactory system also needs to detect and identify odors in the presence of background mixtures. To probe this ability, we tested sFlt1 mice on an olfactory figure-background segmentation task (Rokni et al., 2014; Lebovich et al., 2021). Mice were trained to perform a task, in which they were required to report whether presented mixtures contained a target odorant (see methods). The difficulty of the task was gradually adjusted by presenting mixtures that are composed of more components (Figure 2i). All 8 sFlt1 mice performed this task above chance level (Figure 2j). Although, we did not find a statistically significant difference in the performance of control and sFlt1 mice, the difference in performance between individual sFlt1 mice was to a large extent explained by OB size (r=0.85, p=0.008, Supp Figure 2). These data show that despite the apparent similarity in behavioral performance, more challenging tasks may reveal the deficits associated with OB degeneration.

### Odors drive normal cortical responses in sFlt1 mice

The integration of apparently randomly organized OB projections gives rise to a distributed code for odors in the piriform cortex (PCx) that is thought to underlie odor recognition (Illig and Haberly, 2003; Rennaker et al., 2007; Poo and Isaacson, 2009; Stettler and Axel, 2009; Gottfried, 2010; Isaacson, 2010; Wilson and Sullivan, 2011; Miura et al., 2012; Bekkers and Suzuki, 2013; Giessel and Datta, 2014; Roland et al., 2017; Blazing and Franks, 2020). The severe degeneration of the OB is expected to greatly reduce the information content of the input into PCx and therefore also its capacity to encode odors. However, the fact that mice were able to perform the odor-guided tasks suggests that PCx may be able to encode odors despite OB degeneration. To address this question, we recorded neural activity in the PCx of sFlt1 mice. We implanted tetrodes in two mice with 10% and 14% of normal OB volume (Figure 3a) and recorded the activity of 152 well-isolated neurons when mice were awake and passively exposed to odors. We presented these mice with the 8 odors that were used in the target-background detection task and analyzed the statistics of neural responses.

**Figure 3.**
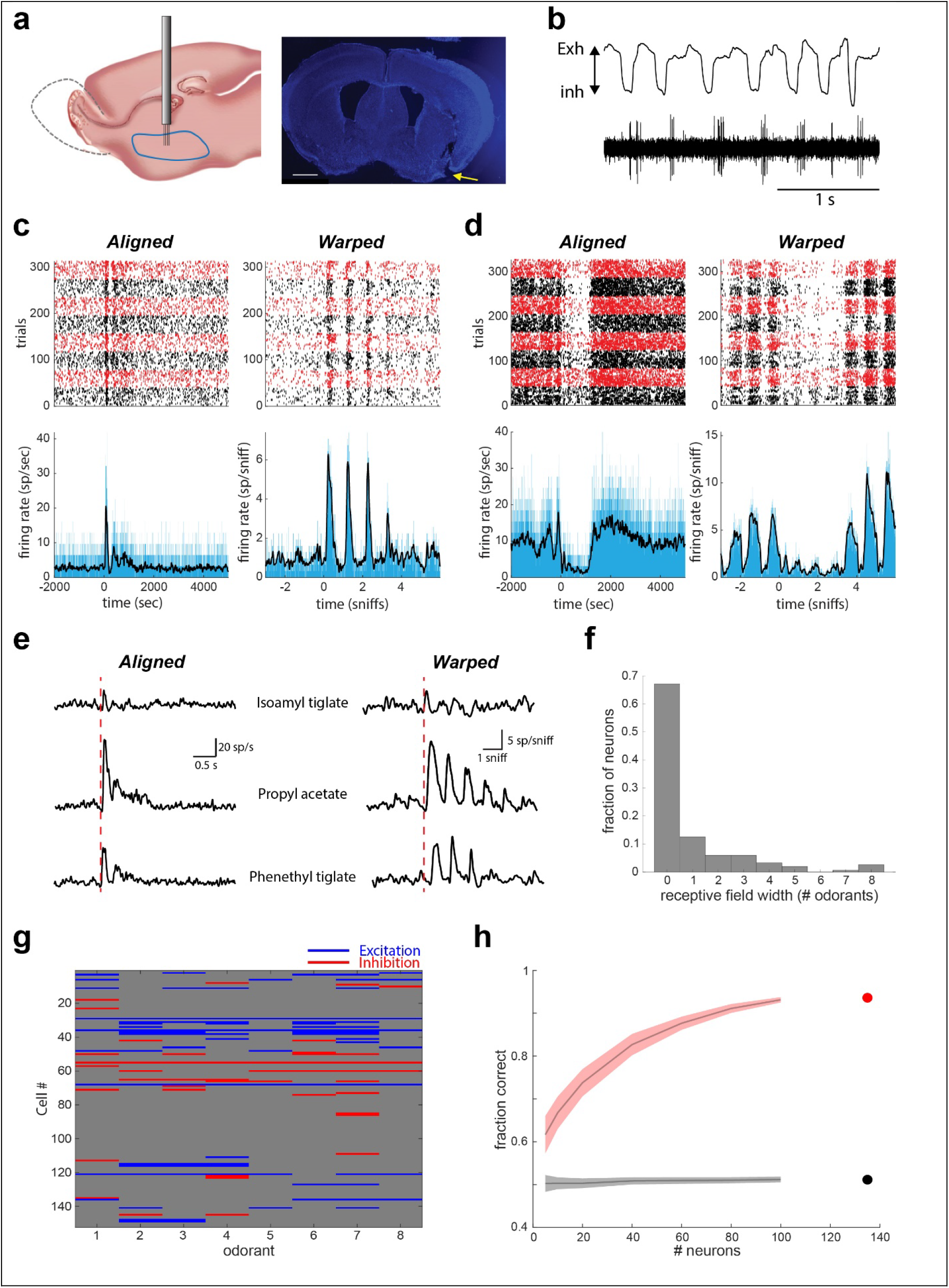
Sniff-locked, odor responses in the aPC. (**a**) A recording tetrode drive (8 tetrodes, 32 channels) was implanted in the aPC. Mice (10% and 14% of normal OB volume, 1 male and 1 female) were presented with 8 different odorants for 1sec with a 9sec interval between trials and underwent >300 trials per session. After each session, the tetrodes were lowered by 75μm. Right: coronal brain section showing the lesion in the aPC made by the implant (arrow). (**b**) Example raw data showing the signal from one wire and the inhalation trace. Note locking of spikes to the breathing cycle. (**c**) Raster plot (top) and PSTH (bottom) for a representative neuron responding to odors by excitation. Time 0 represents the onset of first inhalation after opening of the odor valve. Inhalation-aligned firing is shown on the left and respiration time-warped analysis on the right. The responses to each of the odorants were sorted and each odorant is indicated by alternating black and red blocks. PSTHs show the average response to all odor presentations. (**d**) Same as c, for a neuron responding to odors by inhibition. (**e**) PSTHs of responses to specific odors from the same neuron in c. (**f**) A histogram of tuning width for all recorded cells. Tuning width is defined as the number of odors eliciting a significant response (d’ > 1). (**g**) A summary of the responses of all recorded neurons. Blue and red represent significant excitatory and inhibitory responses (effect size > 1), respectively. Grey represents nonsignificant odor responses. (**i**) Decoding accuracy as a function of the number of neurons used. Decoding was performed using linear classifiers with 10-fold cross validation. 100 random subpopulations of neurons were used for each population size and the mean performance ± SEM is shown. Real data classification is shown in red, and shuffled labels classification in black.

Overall neuronal activity in these sFlt1 mice with less than 15% remaining of the OB, showed similar characteristics to those described in normal rodents (Poo and Isaacson, 2009; Stettler and Axel, 2009; Zhan and Luo, 2010; Miura et al., 2012). First, firing of PCx neurons was modulated by the breathing cycle (Figure 3b-d). Second, individual cells typically responded to a small number of odors (Figure 3e-g). 11% of all cell odor pairs yielded statistically significant responses with individual odors activating 7 to 16 % of the cells (Poo and Isaacson, 2009; Stettler and Axel, 2009; Miura et al., 2012; Bolding and Franks, 2017; Roland et al., 2017). Third, odor responses included both increases and decreases in firing rate (Figure 3c,d,g) (Zhan and Luo, 2010; Bolding and Franks, 2017; Roland et al., 2017). Typically, all responses of an individual neuron were of the same type (either excitation or inhibition, Figure 3g). To describe response statistics without relying on categorization of cell-odor pairs as responsive or non-responsive, we calculated population and lifetime sparseness (Rolls and Tovee, 1995; Willmore and Tolhurst, 2001). Population sparseness of individual odor responses was in line with previous measurements in rodents (mean 0.69 range 0.52-0.84). (Poo and Isaacson, 2009; Miura et al., 2012). Interestingly, lifetime sparseness was lower in the sFlt1 mice (mean 0.29, range 0.02-0.77) (Poo and Isaacson, 2009; Miura et al., 2012; Roland et al., 2017). This may either reflect a decrease in the sparseness of the input into the cortex (for instance if mitral cells integrate input from several receptors), or changes in the local circuitry within PCx. These data show that PCx responses in mice with degenerated OBs show remarkably similar statistics to those that have been previously described in normal rodents.

To guide behavior, PCx responses must be informative about odor identity. To test whether indeed neural responses carry information about the identity of the presented odors, we pooled neurons across mice and decoded the odor identity using linear classifiers (Figure 3h). All 8 odors were decoded with an accuracy of over 85%. The dependence of decoding accuracy on the size of the neuronal population was similar to previously described decoding in healthy rodents (Miura et al., 2012; Bolding and Franks, 2017). In summary, the analysis of neural recordings from PCx in sFlt1 mice with less than 15% remaining of the OB, clearly demonstrate that despite the degeneration of its main sensory input source, PCx receives odor information and encodes odor identity.

### Aberrant circuitry underlies olfaction in sFlt1 mice

Several features of the stereotyped circuitry upstream from piriform cortex are thought to be essential for olfaction. Specifically, odors are first encoded by a large number of olfactory receptors (ORs) in the OE, and projected to the OB. There, the signals from thousands of sensory neurons expressing the same OR are integrated by a few tens of mitral and tufted (M/T) cells within glomeruli. This summation provides a high signal-to-noise ratio in the OB output. Specific layering within the OB provides the anatomical substrate for odor processing within the OB. What circuit may underly olfaction in mice with degenerated OBs? To answer this question, we analyzed the anatomical and transcriptional landscape of OSNs as well as the anatomy of MCs in mice with the degenerated OBs.

### Olfactory sensory neurons

Olfactory processing begins with the specific binding of odorants to olfactory receptors expressed by OSNs. We first asked whether any changes are evident in the OE of sFlt1 mice. To analyze OSN morphology, we immunostained sections of the OE for the OSN marker Olfactory Marker Protein (OMP, (Farbman and Margolis, 1980)). We found that the OE is heavily populated by OSNs that show normal morphology, namely a major sensory dendrite extending from the soma and ending in a ciliary bouton in the nasal cavity (Figure 4a-b).

**Figure 4.**
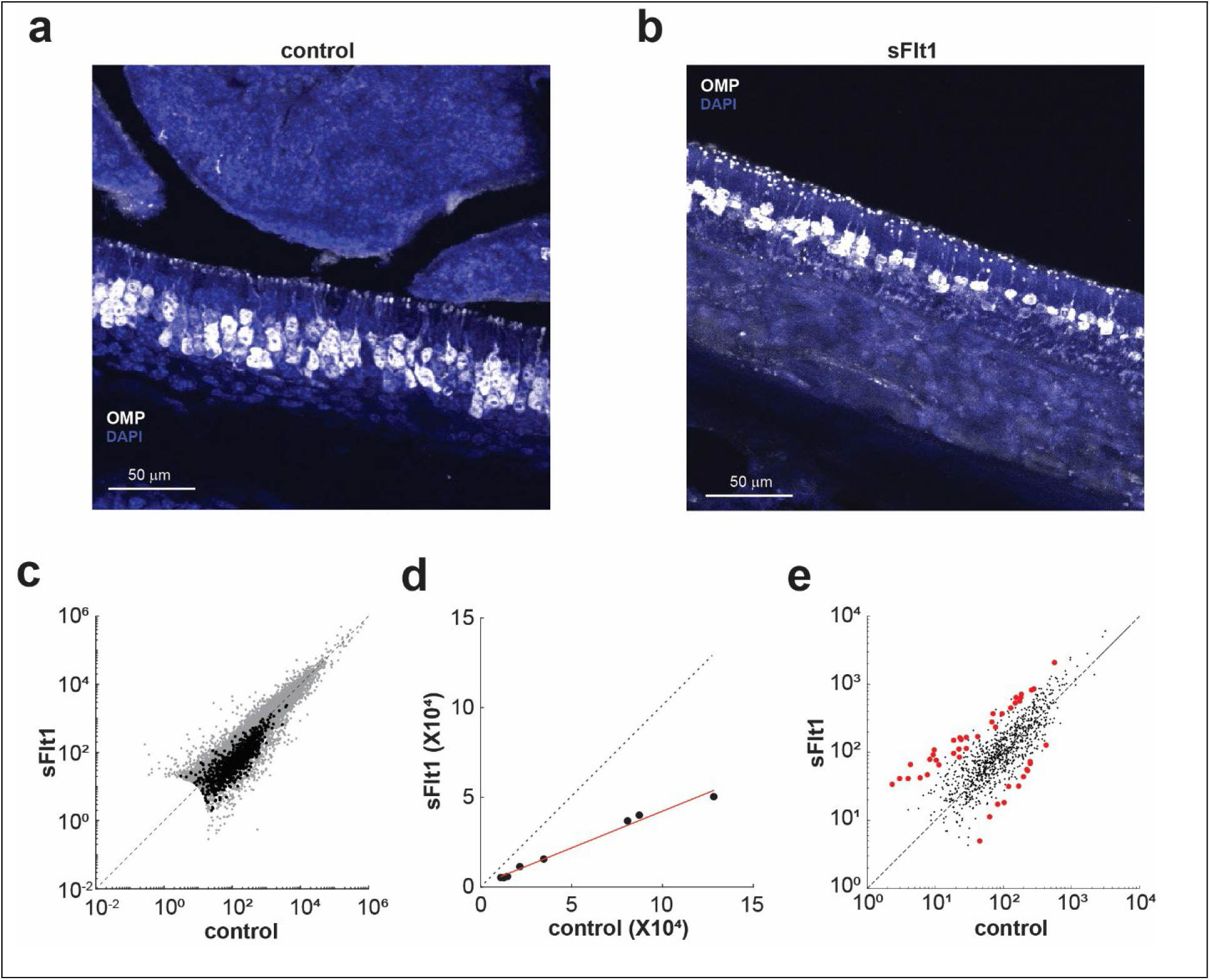
OSN morphology and transcriptomics in sFlt1 mice. (**a-b**) OMP immunostaining reveals no evident differences in cellular morphology of OSNs in control (**a**) and sFlt1 (**b**) mice. (**c**) RNAseq analysis of the olfactory epithelium. The average expression level (normalized reads) of each gene in sFlt1 mice (n=3) is shown as a function of the average expression level in controls (n=3 mice). Olfactory receptor genes are shown in black and other genes in gray. (**d**) Average expression levels of 8 OSN-specific genes (Omp, Cnga2, Adcy3, Ano2, Gnal, Gng13, Stoml3, Cngb1) in sFlt1 and control mice. Red line is a linear fit (slope of 0.41). (**e**) Expression of olfactory receptors normalized in each animal to the level of OSN-specific genes. Red dots indicate significant difference between sFlt1 and control mice.

Mice have over a thousand olfactory receptor (OR) genes (Buck and Axel, 1991). We asked whether the reduced volume of the OB affects the number and expression profiles of ORs. To test this, we performed bulk RNAseq analysis of the whole OE in 3 sFlt1 mice (11, 28, and 29% of normal OB size) and 3 control mice. We found that despite the degeneration of the OB, sFlt mice express the full repertoire of ORs (93% of the ORs that were detected in the controls were also detected in the mutants and 98% of the ORs that were detected in the mutants were also detected in the controls). Comparing OR RNA levels between sFlt1 and control mice, we found that OR expression in the mutants was highly correlated with controls (r=0.89, p<10^−10^), yet also generally reduced (Figure 4c). The slope of a linear fit between the control and mutant expression levels was 0.54. Analysis of non-OR genes that are specific to OSNs (Omp, Cnga2, Adcy3, Ano2, Gnal, Gng13, Stoml3 and Cngb1) (Khan et al., 2013; Ibarra-Soria et al., 2017) revealed lower counts of these genes in the mutant tissue as well (Figure 4d). We interpreted the decrease in these OSN-specific genes as reflecting a reduction in the number of OSNs and normalized all OR counts to these genes (see methods). On average the OSN-normalized receptor counts in sFlt1 mice were 44% higher than those in control mice (p<10^−10^, Wilcoxon singed rank test), suggesting that the expression level of receptor mRNA per cell was higher in the mutants. Further analysis of the relative OR expression retrieved 53 OR genes whose expression is significantly changed in the mutant. 41 ORs were upregulated in the sFlt1 mice and 12 were down-regulated (Figure 4e). We conclude that sFlt1 mice maintain the full repertoire of ORs and that the expression level per cell is increased. This increase may compensate for a reduction in the number of OSNs.

We next analyzed the projections of OSNs within the OB by immunostaining for the olfactory marker protein (OMP) in brain sections. In the normal OB, OSN axons populate the glomerular layer (Figure 5a). In sFlt1 mice the organization of the OSN projections within the OB was strongly dependent on the extent of OB degeneration. The more degenerated the OB, the more axons of OSNs penetrated deep into the OB (Figure 5a-b). In the highly degenerated OBs of 10% or less, we found no glomerular layer at all. Instead, OSN axons occupied the full volume of the remaining OB forming a large unorganized mass. Similarly, DAPI staining revealed no organized layering of cells within these extremely degenerated OBs. In moderately degenerated OBs (20-30 %), the outer layers contained an abnormally dense and unorganized mass of OSN axons, and glomerular-like structures were sometimes evident within the deeper layers (Figure 5b-c). Abnormally large glomeruli were occasionally encountered in these moderately degenerated OBs, possibly reflecting fusing of multiple glomeruli (Figure 5c). To test whether OSNs expressing the same OR retain normal convergence in the OB despite the apparent lack of glomerular organization, we crossed the M72-ChR2-GFP mouse line with the sFlt1 transgenic system. While we could easily locate the M72 glomerulus in control mice (Supp Figure 5), we could never find this glomerulus in sFlt1 mice. M72 OSNs were clearly evident in the OE of both control and sFlt1 mice as were their axons within the nerve layer entering the OB. However, High levels of autofluorescence in the necrotic OB tissue, prevented identification of M72 axons within the OB. These results indicate that OSN axons expressing the same OR do not converge in the degenerated OB.

**Figure 5.**
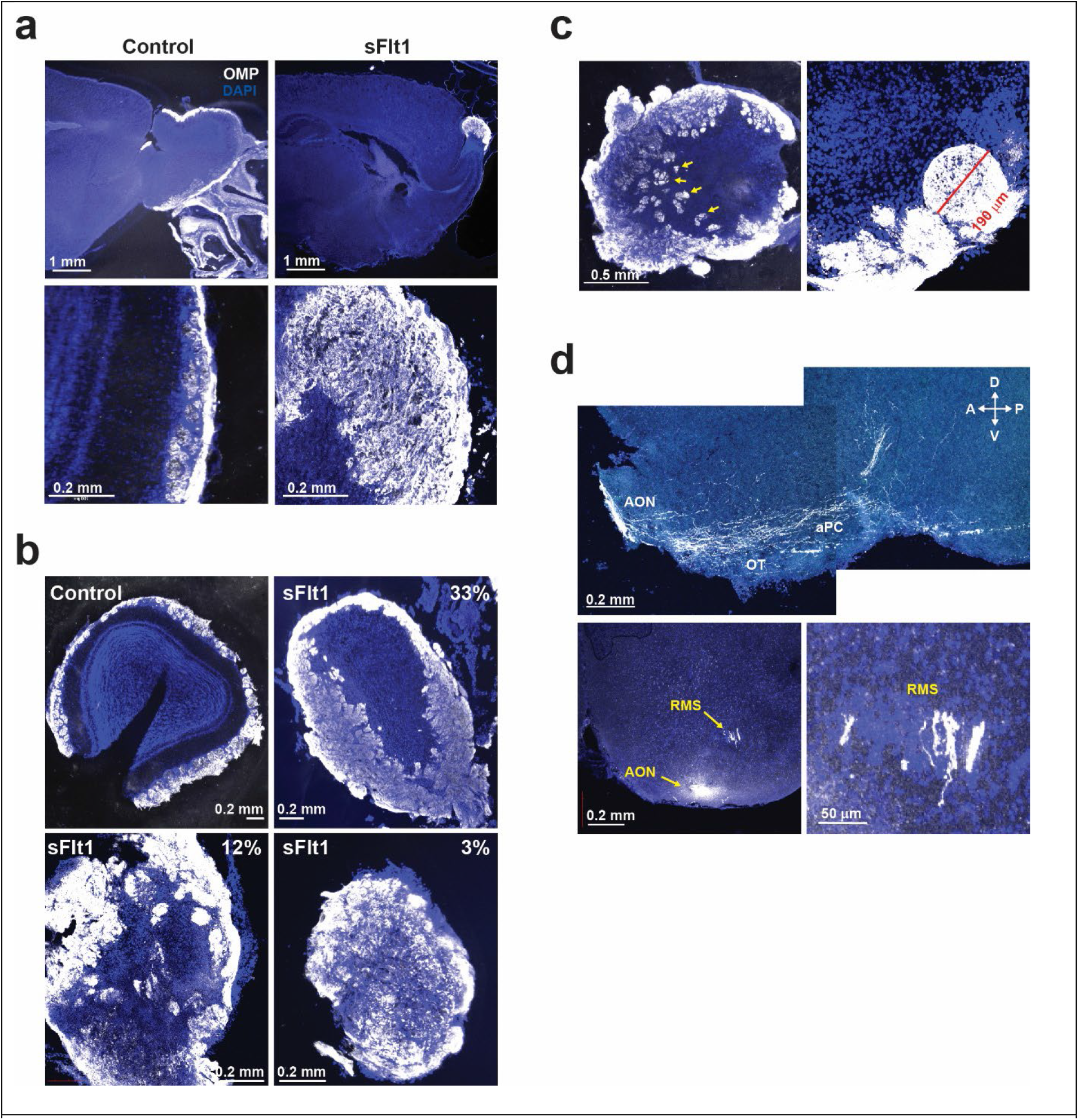
OSN projections in sFlt1 mice. OMP immunostaining. (**a**) Sagittal sections from a control mouse (left) and from a sFlt1 mouse with 10% of the OB (right) showing loss of glomerular structure in the sFlt1 degenerated OB. This mouse participated in the experiment shown in Figure 2f. (**b**) OMP-stained sections of 1 control mouse and 3 sFlt1 mice with 33, 12, and 3% remaining of the OB. (**c**) Sections of the OB of a sFlt1 mouse with 20% OB showing ectopic glomeruli in the center of the OB (arrows), and abnormally-large glomeruli (right). (**d**) OMP+ axons in the AON, the olfactory tubercle, and anterior piriform cortex, as well as in the RMS.

Interestingly, ectopic OSN axons were also found outside the OB in the anterior olfactory nucleus (AON), piriform cortex, the olfactory tubercle, and the rostral migratory stream (RMS) (Figure 5d). Projections of OSN axons to some of these abnormal places (i.e. AON and RMS) were also seen in rats that were bulbectomized at a young age (Slotnick et al., 2004). The fact that ectopic OSN projections are predominantly seen in olfactory brain regions despite their distance from the OB is intriguing. It should be noted however that only a small minority of OSN axons projected to these olfactory cortical areas. The vast majority of OSNs still projected to the degenerated OB.

### Mitral/tufted cells

We next analyzed the organization M/T cells in the rudimentary OB. In order to visualize M/T cells, we created a quadruple transgenic mouse by crossing sFlt1 mice with the Tbet-Cre line (Haddad et al., 2013), and with a Cre-dependent TdTomato reporter line (ai9). The whole mount view of the OB (Figure 6a) revealed presence of M/T cells in the OB and their projection to the olfactory cortex via the lateral olfactory tract (LOT). We analyzed the organization of M/T cells in coronal slices of the OB. The density of M/T cells in sFlt1 mice was not significantly different from their density in controls (cells/mm^3^ counted in 50μm-thick slices), indicating that the reduction in the total number of M/T cells was proportional to the reduction in OB volume (Figure 6b). Generally, the organization of M/T cells and its dependence on the severity of degeneration matched the organization of OSN projection with in the OB. As OB degeneration increased, the M/T cell layer became less organized and eventually could not be defined in the most severely degenerated OBs (Figure 6c,d). To test whether OSN projections within the depth of the OB may have postsynaptic counterparts, we stained slices from sFlt1-Tbet-TdTomato animals with antibodies for OMP. We found that most regions innervated by OSN axons (including ectopic glomeruli) are also populated by M/T cell processes, potentially forming synaptic connections there (Figure 6d). To better visualize the apical dendritic arbors of individual M/T cells and analyze their morphology, we sparsely labelled them with viral injections of retrograde AAV expressing mCherry into the aPC. As expected, we found that M/T cells have abnormal morphology in sFlt1 mice. While in control animals all M/T cells had a dense dendritic arbor limited to a single glomerulus and a soma located in the vicinity of this glomerulus (Figure 6e, top), apical dendrites in the sFlt1 mice spread over larger distances and seemed to occupy the space of several putative glomeruli without forming dense tufts (Figure 6e, bottom). Additionally, in some examples, the soma was not located directly beneath the apical dendrite but rather far to the side (Figure 6e, right). We next analyzed M/T cell projections, namely the LOT. We mounted coronal slices at the level of the anterior and posterior piriform cortex. Although the LOT was found in its normal location, its volume was dramatically reduced, as expected from the major decrease in the numbers of M/T cells (Figure 6f). Overall, these data show that highly abnormal OB organization underlies olfaction in sFlt1 mice.

**Figure 6.**
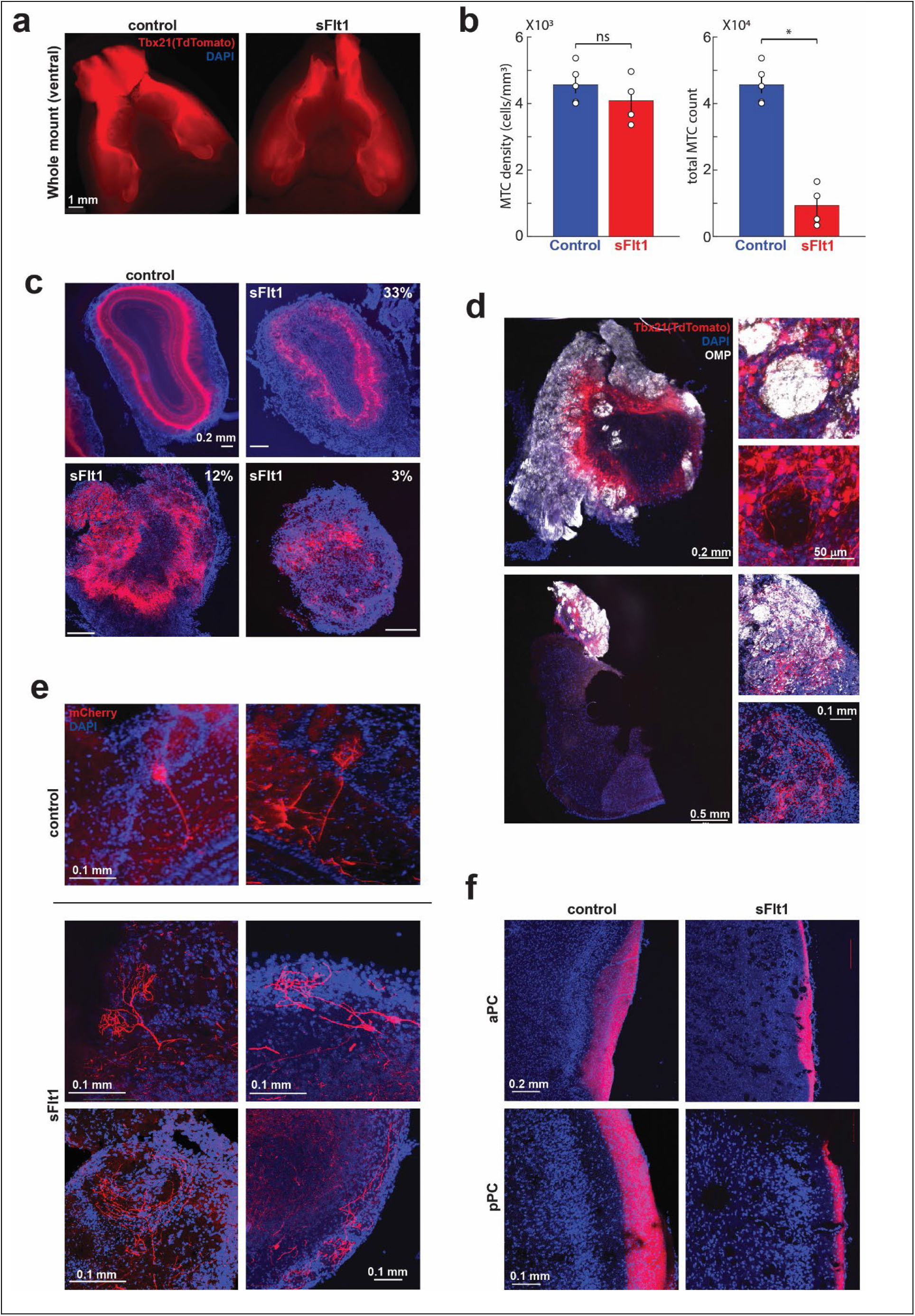
M/T cell organization in sFlt1 mice. The Tbet-cre mouse line and the Ai9 line (TdTomato reporter) were crossed to the sFlt1 system to visualize M/T cells. (**a**) Whole mount view of the ventral side of the brain showing the M/T axonal tract (the LOT). (**b**) Quantification of the total M/T cell numbers in one OB (P=0.0159, Wilcoxon rank sum test) and of the density of cells (P=0.322). N=5 control, 4 sFlt1. (**c**). Slices of mice with gradual loss of OB tissue showing disorganized Tbet+ cells in in the sFlt1 mouse. Recordings from the PCx of the mouse with 14% are shown in Figure 3. (**d**) Immunostaining for OMP reveals that the disorganized OSNs nerve endings and ectopic glomeruli in the sFlt1 mouse contain processes of Tbet+ cells (arrows). Two representative examples are shown. (**e**) A retrograde AAV virus expressing mCherry was injected to the aPC. Slices from the OB show individual M/T cells. Two representative examples for M/T cells from a control mouse and 4 from a sFlt1 mouse are shown. Note abnormal morphology of M/T apical dendrite expanding beyond the territory of a single glomerulus. (**f**) Images of the anterior and posterior piriform cortex showing reduced thickness of the LOT in the sFlt1 mouse.

## Discussion

We used a mouse model with developmental OB degeneration to study how olfactory functionality remains largely intact despite an almost absent OB. The use of a developmental model for OB degeneration allowed us to explore the full capacity of compensatory plasticity mechanisms and to describe in detail the histological structure that is formed in the remaining OB. Some of our findings are in line with previous studies that removed the OB surgically (Lu and Slotnick, 1998; Slotnick et al., 2004; Erskine et al., 2019). We show that mice with as little as a few percent of the OB have a remarkably functional sense of smell. Not only can these mice perform odor-guided behaviors as assessed by structured behavioral tasks, they also maintain the innate valence of natural odors. Connectivity between the OE and the brain in these mice is mostly maintained via the residual OB, but also by direct projections of OSN axons to cortical regions. Although the anatomy of the remaining OB was highly disturbed, with disordered spatial organization and a great reduction in the number of principal neurons, piriform cortex maintained almost normal odor responses that could underly behavioral performance. The sFlt1 mouse is therefore a great model for studying how sensory function is maintained despite extremely aberrant underlying circuitry.

Is the OB so redundant that only a small portion of it suffices to support normal function? The answer to this question is probably no. We think that it is important to keep in mind that behavioral tests of olfaction in the lab are far less demanding than the use of this sense in real life and natural environments. The need to identify and localize odors with varying concentrations and plume dynamics on top of rich and dynamic backgrounds is hard to mimic in the lab. Our finding that the performance of the olfactory figure-background segmentation task depends on the volume of the remaining OB supports this view. It will be important in the future to develop novel tasks that capture more of the real-world complexities of the olfactory sensory scene.

We find that basic odor response properties in PCx are maintained in mice with ~10% of the OB. The number of odors activating individual neurons and the number of neurons activated by individual odors are both similar to reports in normal mice and rats (Poo and Isaacson, 2009; Stettler and Axel, 2009; Miura et al., 2012; Roland et al., 2017). Additionally, the number of neurons required for decoding odor identity is similar to previous reports (Miura et al., 2012; Bolding and Franks, 2017). These properties of piriform cortical responses are in line with distributed coding of odor identity that has been previously proposed (Haberly, 2001; Isaacson, 2010; Giessel and Datta, 2014; Blazing and Franks, 2020). Interestingly, we found that the average lifetime sparseness in mice with degenerated OBs was lower than reports in normal rodents (Poo and Isaacson, 2009; Miura et al., 2012; Roland et al., 2017). This may reflect M/T cells integrating inputs from several ORs. This integration is expected to decrease the lifetime sparseness of M/T cells and therefore also the lifetime sparseness of neurons in PCx.

Beyond the lower lifetime sparseness in PCx, two other findings suggest that M/T cells in sFlt1 mice integrate input from OSNs expressing different ORs. First, we found no evidence for convergence of the M72 OSN axons in the OB. Second, we found that M/T cell apical dendrites cover abnormally large regions of the OB. While this may simply reflect missing guiding signals in the degenerated OB (Sakano, 2010; Mori and Sakano, 2011), we speculate that integration of signals representing multiple ORs compensates for a reduction in the number of mitral cells. As sFlt1 mice maintain the full repertoire of ORs, the number of M/T cells representing each OR will be greatly reduced. M/T cells respond to odors with great trial to trial variability (Rinberg et al., 2006; Fantana et al., 2008) and the great reduction in their numbers is expected to greatly reduce the fidelity of the signals they carry. Forgoing glomerular organization may allow spread of the signal over a larger number of M/T cells and potentially increase its fidelity.

The sFlt1 mouse model recapitulates the incidental cases of humans that have a functioning sense of smell despite no apparent OBs (Rombaux et al., 2007; Weiss et al., 2020), and suggests that these subjects may rely on small degenerated OBs. Additionally, direct projections from the OB to the olfactory cortex may contribute to their ability to smell. We believe that our study not only sheds light on putative compensatory processes but also proposes a mechanism for the pathogenesis of OB degeneration in humans. In our model, the fragility and instability of OB blood vessels at a specific time window during development led to its specific degeneration. Although we inhibited VEGF signaling artificially by overexpressing a “VEGF-trap” molecule, VEGF can also be dysregulated during development in humans, for instance, in preterm babies which are more prone to be affected by oxygen imbalance and may undergo improper oxygen treatment at neonatal intensive care units. Previous studies have shown that retinopathy of prematurity (ROP), a preterm condition leading to blindness, is caused by VEGF imbalance due to hypoxia/hyperoxia episodes and damage to the immature retinal blood vessels (Hartnett, 2015). Not only tissue oxygenation (and fluctuations in VEGF levels) may damage OB vasculature, but also any other vascular impairment during the relevant time window in pregnancy or in the preterm. Interestingly, prematurity or low birth weight was highly correlated with left-handedness (Domellöf et al., 2011; Heikkilä et al., 2018; Marlow et al., 2019) and even specifically in premature girls (Fagard et al., 2021), which corresponds with the high prevalence of left-handed women to lack the OB (Weiss et al., 2020). OB degeneration in humans may therefore potentially also be a syndrome of oxygen imbalance during development.

## Methods

### Mice

All animal procedures were approved by the animal care and use committee of the Hebrew University. Both males and females were used. Transgenic mouse lines used in this study were as follows: the CamkIIα-tTA (Mayford et al., 1996), Ai9 (Madisen et al., 2010), Tbet-cre(Haddad et al., 2013) and M72-GFP(Smear et al., 2013) were purchased from Jackson Laboratories (strains 007004, 007909, 024507, 021206); the pTET-sVEGFR1 responder line was as described previously (Licht et al., 2010). Due to the use of multiple transgenic lines, we performed experiments on a C57/BL6-ICR cross-breed. After the detection of vaginal plugs, the drinking water of pregnant females was supplemented with 500 mg/l tetracycline (Bio Basic Canada, #TB0504) and 3% sucrose. For induction of sFlt1 at embryonic day 13.5, tetracycline-supplemented water was replaced by fresh water. Genotyping was done after weaning. Littermates that inherited CamkIIα-tTA or pTET-sVEGFR1 alone served as controls. Regular specific pathogen–free housing conditions with a 12-hour light/ 12-hour dark cycle were used. Irradiated rodent food and water/tetracycline were given ad libitum.

During behavioral experiments, animals were housed in a reversed light/dark facility and all behavioral training and testing were done during the subjective night time. Water/food consumption was restricted according to testing.

### OB volume measurements

Most OB volumes were calculated with magnetic resonance imaging (MRI). In some cases, fixed brain slices were used to estimate volume. MRI experiments were performed at the Wohl Institute for Translational Medicine at Hadassah Hebrew University Medical Center. MRI images were acquired on a 7T 24 cm bore, cryogen-free MR scanner based on the proprietary dry magnet technology (MR Solutions, Guildford, UK) using 20-mm inner diameter, 18 mm length mouse brain quadrature volume coil. For MRI acquisition, mice were anesthetized with isoflurane vaporized with O_2_. Isoflurane was used at 3.0% for induction and at 1.0 - 2.0% for maintenance. The mice were positioned on a heated bed, which allowed for continuous anesthesia and breathing rate monitoring throughout the entire scan period. Coronal, axial and sagittal T2-weighted (T2W) spin-echo images were acquired for anatomical segmentation purposes (repetition time = 3000 milliseconds, echo time = 45 milliseconds, FOV= 20mm, in-plane resolution = 78 μm, slice thickness = 0.4 mm). Volume calculation was done using ImageJ software (NIH). The OB area in each frame was measured manually and the sum was multiplied by the slice thickness (0.4mm).

### Buried food test

Animals were food-deprived for 18h prior to experiment. The chamber consisted of a clean standard plastic cage (44 × 34 × 20 cm) with a 4 cm layer of new bedding. A food pellet (standard mouse chow) was buried in the bedding, approximately 3 cm beneath the surface, at a random location. The bedding surface was smoothed out and the mouse was introduced into the cage. The time spent to locate the buried food was recorded. The maximum test time allowed was 480s. Each animal underwent the experiment 4 times (once a week).

### Attraction/avoidance responses to odorants

Urine was collected at the day of experiment from a male rat or 5-6 female mice. An odorant (urine/chemicals) was smeared on a standard mouse cage’s wall at one random corner. Mice were placed individually in the center of the cage and their behavior was video recorded for 5 min. The percentage of time spent in each of the 4 quadrants of the cage was automatically calculated using a homemade MatLab script. TMT (10μl) was purchased from PheroTec, Delta, Canada. For the female urine experiment, only males were tested.

### Odor discrimination in the ‘Educage’

Training was done in the EduCage, an automated custom-made home cage setting integrated with RFID antenna, that was previously described (Maor et al., 2019). The setting includes a home cage connected through a short tunnel to a small Plexiglas chamber (10×10×10 cm) where behavioral training takes place. Mice are individually identified and trained using an RFIDs (Trovan) that are surgically implanted under the scuff.

Food was provided ad libitum while access to water was only in the EduCage. Mice could engage with the behavioral apparatus and retrieve water without restriction. At the beginning of each experiment, RFID-tagged mice were placed in a large home cage that was connected to the EduCage. Mice were free to explore the EduCage and drink the water at the behavioral port for 24 hrs. Any entry to the port in this habituation stage immediately resulted in a drop of water, but no odor was delivered. Mice progressed through several stages of training. First, each port entry initiated a trial in which the target odor Limonene R (62118 SIGMA) was delivered for 2 seconds and mice had to lick the water spout to retrieve a water reward. Second, target and non-target odors were presented with equal probabilities and mice had to lick only following the target odor. Pinene R, (Sigma-Aldrich W290238) was first used as a non-target odor (easy discrimination), and S-Limonene (Sigma-Aldrich 62128) - the target’s enantiomer - was used in later stages (harder discrimination). Hits were rewarded with a drop of water and false alarms were punished by a mild air puff and a 9 second timeout. 3 sFlt1 mice with >30% of the OB (1 female and 2 males) and 4 littermate controls (2 females, 2 males) were tested.

### Background segmentation behavioral assay

#### Surgery

Mice were anesthetized (Isoflurane), the skull was exposed and a metal head-plate was attached with Metabond^®^ for subsequent head restraining. Mice were allowed to recover for 1 week before water restriction and behavioral training.

<u>The behavioral apparatus</u> consisted of a head restraining device, an odor delivery system, a lick detector and a water delivery system (Rokni et al., 2014) (Figure 2f). Odor delivery, monitoring of licking and water rewards were controlled using computer interface hardware (National Instruments) and custom LabView software. The mouse was continuously monitored under low intensity red illumination using a CCD camera during behavioral sessions. Odorant mixtures were presented using a custom-made 8-odorant olfactometer that was designed to allow the production of any odorant combination without mutual dilution(Rokni et al., 2014). Each module was made of two glass tubes, one containing the odor and solvent and one containing only the solvent. When an odor was off air flowed through the solvent tube and when it was air flowed through the odorant tube. This design yields a constant continuous flow with varying composition. 3-way solenoids (the Lee Company) were used to switch from the solvent tube to the odorant tube. Odorants (Sigma Aldrich) were as follows (CAS number in parenthesis): Ethyl propionate (105-37-3), Isoamyl tiglate (41519-18-0), 1-Ethyl hexyl tiglate (94133-2-3), Citral (5392-40-5), Propyl acetate (109-60-4), Hexanal (66-25-1), 2-Ethylhexanal (123-05-7), Phenethyl tiglate (55719-85-2).

#### Training

Mice were water deprived and only given water as a positive reinforcement during behavioral sessions. Behavioral training started by 1-2 days of free exploration of the behavioral apparatus, followed by 1-2 days of head-restrained lick training of the waterspout (which released water only upon licking). Target-odor detection training began once mice were readily licking the waterspout. Every mouse was arbitrarily assigned a target odor. Mice were presented with pseudorandom mixtures of odorants and were required to lick the water spout if the mixture contained the target odorant (go trial) and refrain from licking if it didn’t (no-go trial). Hits were rewarded with a 3 μL drop of water. For training purposes, the maximal number of odorants in a mixture was limited on early sessions to make the task easier. A total number of 8 sFlt1 mice (males and females), all with <35% of the OB, and 5 control mice were trained on this task.

### Electrophysiology in aPC

#### Tetrode drive surgery

Mice were first anaesthetized using a combination of an i.p. injection of ketamine (50 mg/kg) and medetomidine (10mg/kg) and maintained under anesthetia by isoflurane inhalation. Mice were placed in a stereotactic device (David Kopf Instruments, California, USA) and an incision was made in the skin to expose the skull. Two drill holes (~1mm) were made into the skull: one above the right aPC (coordinates AP 1.5, ML 2.8), and the other on the opposite side for a ground wire. A metal head-plate was then attached to the posterior part of the skull using dental cement (Metabond^®^). A custom-made 3d-printed drive housing 8 tetrodes (platinum/iridium) and a single screw to advance them all was then placed above the aPC cranial window, lowered to its initial position 3 mm deep, and fixed with additional application of dental cement. The ground wire was then inserted into the brain and cemented. A plastic cone with a retractable lid was placed above the implant and fixed into place by clear epoxy resin. The mouse was woken up using an intraperitoneal injection of atipamezole (2 mg/kg) and allowed to recover for 1 week before recording. Two animals (a female and a male) with 10% and 13% of the average normal OB size were used.

#### Recording sessions

Awake mice were head-restrained in the same apparatus used for the figure-background task. Eight single odorants (no mixtures) were presented in random order for 3 s, 10 s apart. Odors were presented through a mask that was also connected to an air flow sensor (Honeywell AWM3300V) for monitoring of breathing. Each session had over 300 trials. Signals from the tetrodes were amplified, sampled at 32 KHz, and digitized with 16-bit resolution (Digital Lynx SX, Neuralynx). Spikes were detected online and later sorted offline using MClust (David Redish). Only clusters with less than 1 in 1000 intervals below 2 ms were considered as single units. Tetrode array placement was validated postmortem.

#### Data analysis

Spike sorting was performed using MClust4.4 (David Redish). Analysis of single unit activity was then performed using custom-written Matlab codes. Statistical significance of odor responses was assessed by comparing the distributions of firing rates in the first sniff of odor presentation vs a control sniff (outside odor presentation), using Cohen’s d effect size. Effect size above 1 was considered significant. Population and lifetime sparseness were calculated as in (Miura et al., 2012) using the following formula:

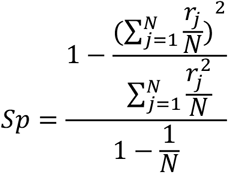

For population sparseness, N is the number of neurons and *r_j_* is the response of neuron j. Responses were calculated as the absolute value of Z-scored firing rates within the first two sniffs. Z-scoring was achieved by subtracting the mean no-odor firing and normalizing by the standard deviation. Lifetime sparseness was calculated using the same formula but j corresponding to the odors and N being the number of odors.

Odor decoding was performed using cells pooled across sessions and mice. 10000 population trials were generated for each of the 8 odors by randomly selecting 1 trial from each of the 152 cells. These trials were classified using the Matlab function fitclinear, with a logistic function learner, sample weighting, and 10-fold cross validation. Chance level performance was evaluated by shuffling the odor labels. The effect of population size on decoding success rate was evaluated by randomly selecting subpopulations of 5, 10, 20, 40, 60, 80, and 100 neurons and repeating the decoding process above. Subpopulations of each size were randomly sampled 100 times and the mean performance was reported.

##### RNA sequencing of OE

OE from 3 controls and 3 sFlt1 mice was dissected and immediately placed in RNA lysis buffer (Zymo Research R1051) without vortexing for 5min and further processed according to manufacturer’s guidelines. ~200ng of RNA was used for the library construction. RNA ScreenTape kit (catalog #5067-5576; Agilent Technologies, Santa Clara, CA), D1000 ScreenTape kit (catalog #5067-5582; Agilent Technologies), Qubit^®^ RNA HS Assay kit (catalog # Q32852; Invitrogen, Carlsbad, CA) and Qubit^®^ DNA HS Assay kit (catalog #32854; Invitrogen) were used for each specific step purpose for quality control of RNA libraries. KAPA Stranded mRNA-Seq Kit with mRNA Capture Beads (Kapabiosystems, KK8421, https://www.kapabiosystems.com/) was used. Library was eluted in 20 ul of elution buffer. All DNA samples libraries were pooled in one tube in equal molarity. Multiplex samples Pool (1.65 pM including PhiX 1%) was loaded in NextSeq 500/550 High Output v2 kit (75 cycles) cartridge (catalog #FC-404-1005; Illumina, San Diego, CA) and loaded on NextSeq 500 System machine (Illumina), with 75 cycles and single-Read sequencing conditions.

##### Analysis of RNA-Seq data

Raw reads were quality-trimmed with cutadapt, v3.5, then poly-G and adapter sequences were removed from the 3’ end. Processed reads were aligned to the mouse GRCm39 genome with Tophat, v2.2.1, allowing for 5 mismatches, using gene annotations from Ensembl release 106. Raw counts per gene per sample were obtained with htseq-count, v2.0.1, then normalization was done with the DESeq2 package, v1.26.0. Noticing a general reduction in the expression of olfactory receptors genes, as well as OSN marker genes, led to a second normalization of olfactory receptors signal, aiming to compensate for the probably smaller population of OSNs in the sFlt1 mice. Following the method described in (Khan et al., 2013; Ibarra-Soria et al., 2017), 8 OSN marker genes were used for a second normalization step. The markers were ENSMUSG00000005864 (Cnga2), ENSMUSG00000020654 (Adcy3), ENSMUSG00000024524 (Gnal), ENSMUSG00000025739 (Gng13), ENSMUSG00000027744 (Stoml3), ENSMUSG00000031789 (Cngb1), ENSMUSG00000038115 (Ano2) and ENSMUSG00000074006 (Omp). The geometric mean of the 8 marker genes was calculated for each sample. The average of those was divided by the individual geometric mean from each sample to give the normalization factor for that sample. All olfactory receptor genes normalized signals were multiplied by this factor and rounded to the nearest integer, then served as input counts for a new DESeq2 analysis without further normalization. Analysis was done with default parameters, except not using the independent filtering algorithm. The significance threshold was taken as padj<0.1 (default).

### Stereotaxic AAV injection

Plasmid #26976 was purchased from Addgene and a retrograde AAV (titer 10^−12^) was prepared by the virus core at The Edmond and Lily Safra Center for Brain Sciences (EVCF). Mice were anaesthetized with isoflurane and placed in a stereotactic device (Kopf). A skin incision was made and a drill hole was made over the aPC (coordinates: AP1.5, ML 2.8, DV 4.4 from Bregma). 0.3μl of the virus was injected by a 10μl microinjector (World Precision Instruments). The craniotomy hole was plugged by bone wax and the skin was sutured. Brain was retrieved after 4 weeks.

### Histology and Immunofluorescence

Brains were fixed by immersion in 4% paraformaldehyde on ice for 12h, washed with PBS, embedded in 2% agarose and sectioned to 50-μm coronal/sagittal sections by a vibratome (Leica). For OE sections, tissue was immersed in 30% sucrose for 2 days followed by embedding in Tissue-Tek OCT and sectioning of 50μm slices in a cryostat (Leica). Staining was done as described (Licht et al., 2015) with the following: anti-GFP (1:400; Abcam RRID:AB_305643) anti-OMP (1:500; Abcam RRID:AB_2858281). Alexa 488 antigoat RRID: AB_2336933 and Alexa 647-anti-rabbit PRID:AB_2492288 were from Jackson Immunoresearch (1:400 dilution). Sections were mounted on glass slides and covered by mounting medium containing Dapi (SouthernBiotech).

### Microscopy

Confocal microscopy was done using Olympus FV-1000 on 10X, 20X and 40X objectives and 2 μm distance between confocal z-slices. Low-magnification images and whole mount fluorescence were acquired using Nikon SMZ-25 fluorescent stereoscope on X1 and X2 objectives. Whole-mount brain images were taken by a mobile phone.

### Mitral/Tufted cells quantification

Tbet-cre crossed to Ai9 animals were used. Quantification was done on confocal Z-stack images of 50μm-thick slices of the OB, taken under 20X objective. The MCL was positioned so that it crossed from one corner of the image to the other (as seen in Figure 3a bottom left image and not always possible for sFlt1 animals). The soma of TdTomato^+^ cells was counted manually by a blind experimenter using FV-10 Fluoview viewer (Olympus). Cell density was calculated as the number of cells per a box of 50X635X635μm (and was normalized to cells/mm^3^). Total number of cells was calculated by multiplying cell density by the OB volume (as measured using MRI). N=4 sFlt1 animals, 6 control animals. 4-8 images were taken for every animal.

