## supplemental figures and tables for "Aberrant circuitry underlying olfaction in the face of severe olfactory bulb degeneration"

**Licht et al. Supplementary information**

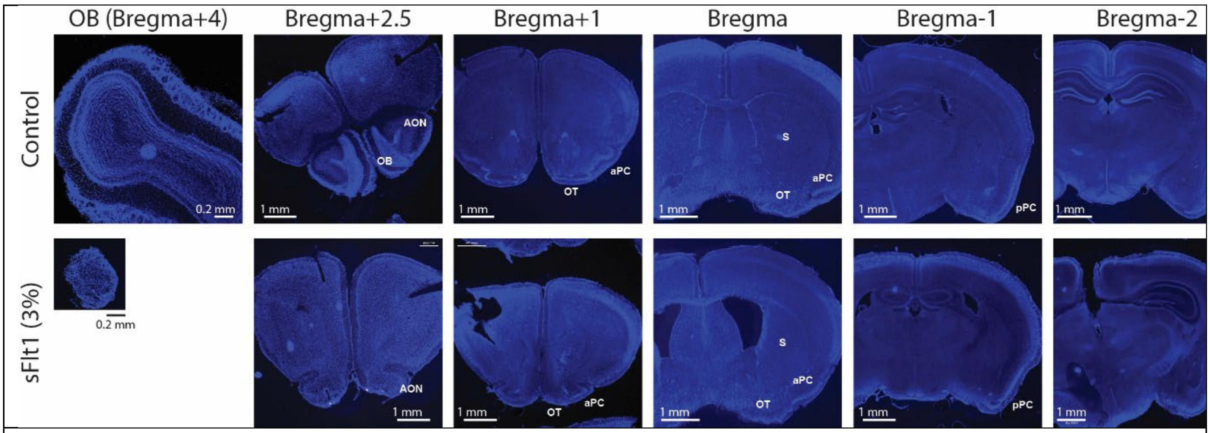


**Supp Figure 1.** Coronal sections of a control (top row) and a sFlt1 mouse with an OB of 3% of normal volume (bottom row), showing the largely normal structure of brain parts that are posterior to the OB. AON- anterior olfactory nucleus; OT- olfactory tubercle; aPC – anterior piriform cortex; pPC – posterior piriform cortex; s- striatum.

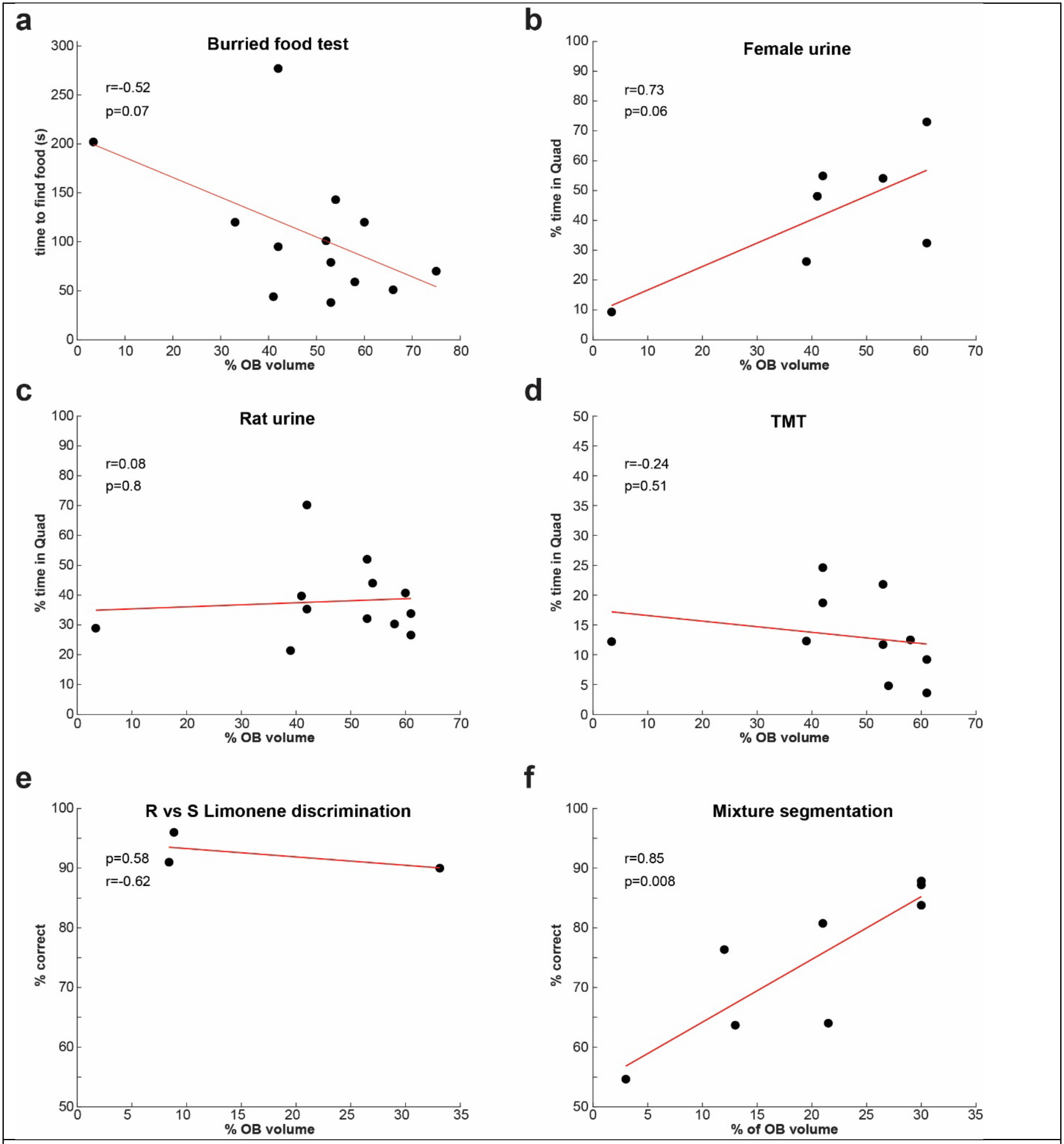


**Supp Figure 2: Behavioral performance as a function of OB size.** (**a**) The time to find buried food. (**b-d**) The time spent in the odor quadrant in an open field test with female urine (**b**), rat urine (**c**), and TMT (**d**). (**e**) Performance on the enantiomer discrimination task. (**f**) Performance on the olfactory figure-background segmentation task. In all plots the correlation coefficients and correlation p-values are shown. Red lines show linear fits to the data.

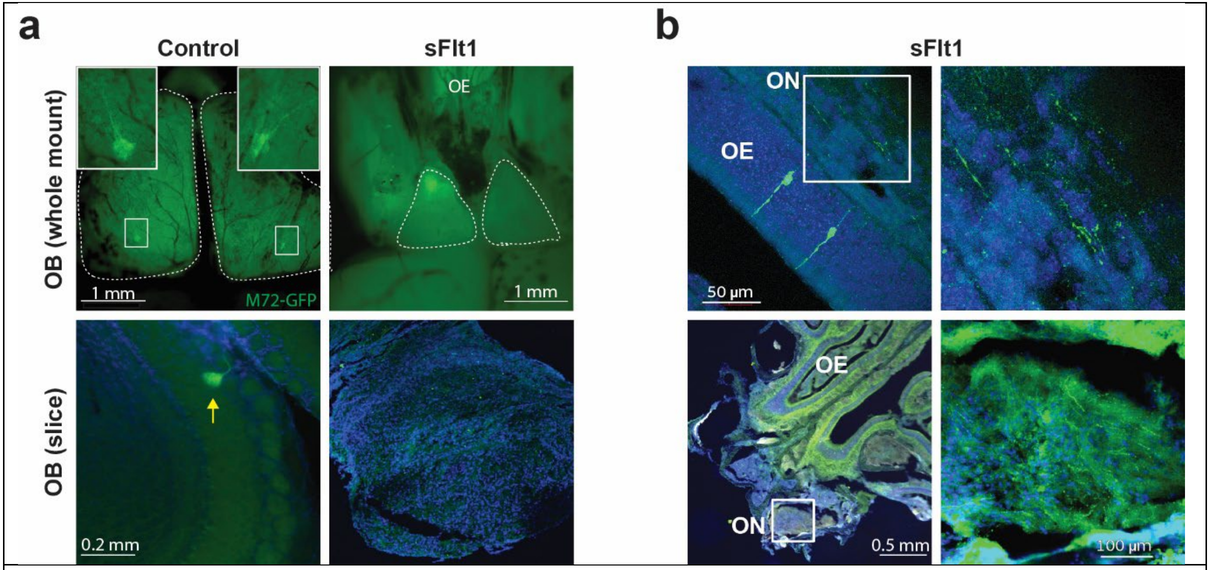


**Supp Figure 5. No evidence for axonal convergence in the OB of sFlt1 mice.** To highlight axons of OSNs of one receptor type, The M72-GFP allele (highlighting OSNs expressing the M72 olfactory receptor) was bred into the sFlt1 system. (**a**) No M72 glomeruli in sFlt1 mice. Whole-mount view of the dorsal surface of the OB (top) and sections (bottom), showing the dorsal M72 glomerulus in a control mouse (arrow) and no evidence for one in a sFlt1 mouse (fluorescence on top left OB is necrotic autofluorescence). (**b**) M72 OSNs and their axons in M72-sFlt1 mice. Top: a section through the OE showing M72 OSNs and M72 axons exiting through the olfactory nerve (enlarged on the right). Bottom: a section through the olfactory nerve entering the OB showing M72 axons there (nerve region enlarged on the right).

**Supp Table 1: Mice used in this study**

| mouse # | sex | vol (mm$^3$) | OB volume (%) | Behavior/recording | Histology in Figure |
|---|---|---|---|---|---|
| 1 | male | 0.84 | 8.4 | Educage | |
| 2 | male | 0.89 | 8.8 | Educage | 5a,d |
| 3 | female | 3.31 | 33 | Educage | |
| 4 | male | 1.39 | 13.8 | Piriform recording | 4b |
| 5 | female | 1 | 10 | Piriform recording | 1c middle, 6d |
| 6 | male | 1.85 | 18.4 | target-background | 6e |
| 7 | male | 3.28 | 32.7 | target-background | Supp5b, 5b and 6c 33%, 6f |
| 8 | female | 3.27 | 33 | target-background | |
| 9 | female | 1.32 | 13.2 | target-background | |
| 10 | male | 1.24 | 12.4 | target-background | 6a, 5b and 6c 12% |
| 11 | male | 2.15 | 21.4 | target-background | |
| 12 | female | 2.10 | 20.9 | target-background | 1c right, 2g, 6e |
| 13 | male | 0.34 | 3.4 | target-background and innate behavior | 1b, 5b and 6c 3%, supplementary 1 |
| 14 | male | 4.13 | 41.1 | innate behavior | |
| 15 | male | 4.21 | 41.9 | innate behavior | |
| 16 | male | 5.34 | 53.2 | innate behavior | |
| 17 | female | 6.04 | 60.1 | innate behavior | |
| 18 | female | 5.34 | 53.1 | innate behavior | |
| 19 | female | 4.25 | 42.2 | innate behavior | |
| 20 | female | 5.77 | 57.4 | innate behavior | |
| 21 | female | 5.40 | 53.7 | innate behavior | |
| 22 | male | 3.87 | 38.5 | innate behavior | |
| 23 | male | 6.10 | 60.7 | innate behavior | |
| 24 | male | 6.08 | 60.4 | innate behavior | |
| 25 | female | 6.56 | 65.2 | cookie test only | |
| 26 | male | 7.53 | 74.9 | cookie test only | |
| 27 | female | 3.31 | 33 | cookie test only | |
| 28 | female | 5.25 | 52.2 | cookie test only | |
| 29 | male | 1.10 | 10.9 | | supp5a |
